# Deer tick virus genotypes are perpetuated by different modes of transmission

**DOI:** 10.64898/2026.03.20.713216

**Authors:** Heidi K. Goethert, Alanna O’Callahan, Richard Johnson, Sam R. Telford

## Abstract

Deer tick virus (DTV), or lineage II Powassan virus, is an emergent tick-borne encephalitis virus in North America. Survivors frequently sustain neurologic sequelae. Nationally reported cases have been increasing. DTV is thought to be maintained in nature by multiple modes including horizontal transmission (from viremic host to tick), cofeeding transmission (between ticks feeding nearby) and by transovarial transmission (female to progeny). Analysis of the relative importance of each mode has been hindered by low enzootic transmission. In 2021, Martha’s Vineyard, Massachusetts experienced an epizootic that allowed us to probe the modes of transmission on the island. We detected virus in 7.8% of questing deer tick nymphs (161 of 2063) and in 0.3% of lone star nymphs (2 of 678). Infected ticks had a highly focal distribution; 56% of infected ticks derived from only 4 of 71 collection sites. Tick mitochondrial genome sequencing demonstrated that infected ticks were not more likely to be siblings than negative ticks and, therefore, were unlikely to have inherited the infection. Whole viral genome sequencing revealed the presence of 3 genotypes, 58% were type1, 0.6% type2, and 13.7% type3. Tick host bloodmeal identification analyses determined that nymphs infected with type1 were significantly associated with having fed on shrews (50 of 94 type1 ticks, odds ratio=2.3, p<0.001). This is consistent with shrews serving as a reservoir. Ticks infected with type3, however, had no host associations, consistent with infection acquired by cofeeding. It may be that local DTV genetic variation is shaped by transmission modes or host associations.

**Importance:** Deer tick virus (DTV; Powassan lineage II) is a tick-borne encephalitis virus that causes a rare zoonosis in North America. Cases have been increasingly reported within the last decade. Is the recent risk trend due to increased transmission? How this virus is perpetuated in nature is not well understood. We took advantage of a natural epizootic on Martha’s Vineyard to probe how the ticks there had become infected. Using a combination of viral whole genome sequencing and bloodmeal remnant identification in ticks, we find that the mode of transmission varied by viral genotype. One genotype is associated with ticks that had fed on shrews, and another did not depend on a specific reservoir host. Host associations may drive genetic diversity of deer tick virus and thus local host population dynamics may influence zoonotic risk.

## Introduction

Powassan virus (POWV) is an emergent tick-borne encephalitis virus in North America. Human infection can lead to disease that ranges from mild flu-like symptoms to severe encephalitis. The case fatality rate has been reported to be as high as 12% (Piantadosi and Solomon, 2022), but this number is likely inflated due to underreporting of mild cases. Surviving patients often experience persistent and debilitating neurological deficits. The index case was a child from Powassan, Ontario who died of encephalitis in 1958 (McLean and Donohue, 1959). Powassan virus has since been identified in the United States, Canada and the far east of Russia. Until the early 2000s, fewer than 35 cases had been reported. Although cases still remain rare, there has been a marked increase in reported cases over the last 20 years: from 2004 to 2016, there were 101 cases (Rosenberg et al., 2018), but just from 2022-2024, 156 cases were reported (CDC, 2026). The basis for an increasing trend is likely due to increased suburbanization and local expansion of deer herds, both of which drive vector tick abundance, as well as an aging human population in such sites, which broadly promotes disease severity. Increased physician awareness and greater availability of testing through state health departments has also enhanced case detection.

There are two genetically distinct POWV lineages, lineage I, referred to as Powassan virus, and lineage II, referred to as deer tick virus (DTV) (Ebel, 2010). For many years, all human cases were attributed to Powassan virus; DTV was thought to be a nonvirulent subtype. In 2009, DTV was definitively identified as the cause of a fatal case of encephalitis (Tavakoli et al., 2009). Although specific POWV lineage generally cannot be inferred for surviving human cases because infection due to these two viruses cannot be distinguished serologically, DTV, not Powassan, has been the cause of more recent cases with definitive molecular sequence identification (Cavanaugh et al., 2017; El Khoury et al., 2013; Piantadosi et al., 2018; Solomon et al., 2018). The two lineages of Powassan virus are thought to be maintained in separate ecological transmission cycles. POWV is maintained among sciurid and medium sized mammals by narrowly host specific *Ixodes* cookei and I. marxi, whereas the vector of DTV is *Ixodes* dammini (the northern clade of *Ixodes scapularis*), hereafter referred to as the deer tick. These tick associations may not be absolute, because deer ticks infected with lineage I Powassan virus have been detected in New York State (Lange et al., 2023). Furthermore, dog ticks (Dermacentor variabilis), lone star ticks (Amblyomma americanum), and longhorned ticks (Haemaphysalis longicornis) are competent vectors in the laboratory (Raney et al., 2022; Sharma et al., 2021). The significance of these other ticks for POWV/DTV risk remains to be determined, although the possibility that lone star ticks may be natural vectors is concerning because they are aggressive human-biters and may be found in some northestern sites where POW/DTV is enzootic (Molaei et al., 2019).

Laboratory and field studies demonstrate that POW/DTV can be maintained by multiple modes of transmission. Larval ticks can acquire infection by feeding on a viremic host and transmit infection after molting to a nymph (transstadial or horizontal transmission). Larvae can also acquire infection by feeding in close proximity to an infected nymph (cofeeding or nonsystemic transmission). Finally, POW/DTV can be inherited by progeny of an infected female tick (transovarial or vertical transmission) (Chernesky, 1969; Costero and Grayson, 1996; McLean et al., 1961; Telford 3rd et al., 1997). Despite these multiple modes of transmission, POW/DTV is rare in ticks, with prevalences < 1% of the ticks in any single site (Anderson and Armstrong, 2012; Brackney et al., 2008; Lange et al., 2024; Xu et al., 2024). The low natural infection rates make it difficult to analyze the enzootic cycle of the virus.

Martha’s Vineyard (MV) is an island off the coast of Massachusetts with hyperendemic Lyme disease and babesiosis. As part of a tick-borne disease prevention program, we have analyzed nymphal deer ticks collected from public walking trails and private yards across the island since 2017. DTV-infected deer ticks are identified every year, but prevalence has typically been less than 1%. In 2021, however, an unusually large number of DTV-infected nymphs were identified, 7.8% of the ticks collected that year. This epizootic allowed us to probe the modes of enzootic transmission of DTV on the island.

We mapped where infected ticks were collected and demonstrate that the distribution on the island is highly focal. We sequenced tick mitochondrial genomes to determine whether infected ticks were related and therefore were potentially the result of transovarial transmission. We used our published retrotransposon method of bloodmeal remnant identification to determine whether the infected nymphs were associated with having fed as larvae on a particular host to distinguish between transstadial transmission and cofeeding. Finally, with the recent invasion of lone Star ticks on the island, we were able to determine whether these ticks contributed to the epizootic. Overall, this epizootic provided a unique opportunity to gain a better understanding of the enzootic cycle of DTV.

## Methods

### Tick collections

Host seeking nymphal deer ticks and lone star ticks were collected by flagging along public walking trails and on private properties on Martha’s Vineyard, Massachusetts as part of their tick-borne infection control program. Supplemental collections occurred in public parks as part of our long-term surveillance efforts on the island.

### Tick processing and virus screening

Ticks were immediately frozen after collection and kept at −20°C until processing. Nymphal ticks were washed with 5% bleach and rinsed twice with PCR-grade water. They were then sorted into individual tubes and homogenized with a sealed p1000 pipette tip along with a few grains of 10 micron zirconium beads. Nucleic acids were extracted using Quick-extract DNA solution (BioSearch Technologies, Hoddesdon, UK). Ticks were incubated in 60µl QuickExtract and then incubated for 6 minutes at 65°C and then 2 minutes at 95°C according to the manufacturer’s instructions. A 5µl aliquot was removed from each tick extract to create pools of 6 ticks. Tick pools were screened for DTV virus using RT-qPCR using previously published primers (El Khoury et al., 2013). Ticks from positive pools were then screened individually to determine which individual was infected.

### DTV mapping and cluster analysis

GPS locations obtained from Google maps for the street address of tick collection sites were used to create maps in ArcGIS (ESRI, Redlands, CA). These locations are not included in the supplemental data to protect the privacy of the property owners. DTV cluster analysis was conducted with SaTScan v10.1 using the discrete poisson model (Kulldorff, 1997).

### Virus quantitation

To estimate the relative viral burden in ticks from our study, positive ticks were reamplified with the DTV screening primers using a duplex qRT-PCR that also amplified tick GAPDH. GAPDH primers and probe were designed using the primer and probe design function within Geneious Prime. Primers are listed in the supplemental primers file. Estimates are expressed as the ratio of Ct from DTV PCR divided by the Ct of the GAPDH PCR. Note that Ct values are inversely related to the amount of presumptive virus in the ticks; so that larger ratios are equivalent to ticks with lower initial copy numbers.

Sequencing and phylogenetic analysis of DTV virus from ticks: The whole viral genomes from PCR-positive ticks were sequenced using a PCR tiling method following a protocol published for COVID virus (Quick et al., 2017). Primers for amplifying overlapping 500bp pieces were designed using PrimalScheme (Grubaugh et al., 2019). Primers are listed in the supplemental primers file. Two PCRs were conducted directly from tick nucleic acid extracts for each sample, one with the odd numbered primers and a second with the even numbered primers. Amplicons were run on a gel to confirm that the PCRs were successful, and then quantitated using a Qubitflex fluorimeter. Individual primers were redesigned using Geneious Prime if amplification of that tile was determined to be less robust than the others. The two PCRs were then combined, barcodes ligated using the Oxford Nanopore Native barcoding kit 24, and then sequenced on a MinION using the R9.1 flowcell, and the newer R10 when it became available, as per manufacturer’s instructions. Guppy was used for basecalling and demultiplexing the barcoded samples using the super high accuracy setting. Low quality reads were trimmed, and primers and barcodes were removed using BBDuk within Geneious Prime. Reads were then mapped to the DTV genome (Genbank HM440559) using minimap within Geneious Prime. Variants with a prevalence greater than 15% were detected, areas of low coverage (less than 15 reads) were masked, and then these changes were mapped onto the reference sequence. Because Oxford Nanopore is notorious for errors in homopolymer strings, all strings 5 bp and longer were manually corrected to that of the reference genome. Assembled genomes were aligned along with 2 genomes from GenBank (HM440559 and OL704186) using MAFFT and neighbor-joining consensus trees were made using Geneious TreeBuilder with 500 bootstrap replicates. Consensus sequences for types 1 and 3 were made by aligning the tick genomes for each type with MAFFT within Geneious. These consensus sequences were then aligned with the sequences from GenBank, translated and the SNPs that resulted in amino acid changes were determined.

### Amplicon genotyping of variants

Due to the relative difficulty of obtaining whole viral genomes directly from field-collected tick extracts, new PCR primers were designed to enable genotyping from sequencing a single small amplicon. Whole genomes from representative genotypes were aligned in GeneiousPrime, and individual SNPs were mapped. Areas with high numbers of unique SNPs were identified and PCR primers were designed to amplify 2 regions of approximately 350bp that contained multiple SNPs that could each be used to identify genotypes. These primers were then used to amplify directly from tick DNA extracts, and the amplicons were sequenced using an Oxford Nanopore MinION flow cell as described above. Both amplicons were attempted for all ticks, but amplification with just one was sufficient to determine genotype. Genome tiling primers DTV13L/R amplifies a piece that overlaps with MVamp1 amplicon and was used to for typing in some samples that did not amplify with the other primers. Primers are listed in the supplemental primer file.

### Estimation of the contribution of transovarial transmission

If the ticks were infected via transovarial transmission, we would expect that infected ticks collected from the same site should derive from a single infected egg batch. Therefore, we compared the mitochondrial genome of infected ticks derived from the same sites to determine their relatedness. Tick mitochondrial genome sequencing was conducted with the same tiling-PCR method detailed above for viral genome sequencing. The tiling primers used are listed in the supplemental primer file. To determine the amount of error inherent in our assay, we sequenced the mitochondrial genomes of 4 larvae derived from a single egg batch from a laboratory colony female. These sequences ranged from 99.95% to 99.99% similar. We, therefore, set 99.95% sequence similarity as our cut-off for assigning two ticks as potentially being from the same egg batch. Using individual collection sites with more than 5 DTV-positive ticks, we then determined the number of ticks that had potential siblings from each site. We did the same from ticks from control sites and compared the outcomes. Control sites were chosen as sites sampled during 2021 with more than 20 deer ticks collected at each site with no detectable DTV. Because these sequences were produced by PCR-tiling, there are instances where the amplification of an individual tile did not yield enough reads, creating gaps in the sequence that are filled by Ns. Geneious treats Ns as being different in a sequence similarity matrix; so sequences were trimmed to remove these areas to get an accurate estimate of similarity. The similarity matrixes are included in the supplemental files.

### Bloodmeal identification

Bloodmeal identification was conducted using our published method that uses qPCR to amplify residual mammalian retrotransposon DNA left in the tick gut from the previous bloodmeal host. All 161 DTV-positive ticks and 217 ticks from 9 control sites were tested. A detailed protocol with primer and probe sequences has been published and recently updated (Goethert, 2026). Briefly, 2µl of tick DNA was used in a 20µl real time PCR using ssofast probe super mix (Bio-Rad Inc. Hercules, CA) for 50 cycles. Ticks were run in duplicate, and any sample which only had a single positive was repeated. Positives that could not be replicated were deemed to be negative. PCRs were run in multiplexes; mouse/vole/rabbit, skunk-raccoon/shrew/deer, and bird/opossum/squirrel-chipmunk were run for all samples. Ticks that were negative for these hosts, were then tested for jumping mouse, mole, rat and cat. Note that the primers are not species or genus specific and are known to amplify other species from the same family. Detailed information on the known cross reactions of the primer sets can be found in the online protocol. Diversity estimates and t-tests were conducted using Past4.09 (Hammer et al., 2001)

### Genbank submissions

All sequence data from this study have been deposited into the NCBI Sequence Reads Archive under Bioproject ID PRJNA1432943.

## Results

### Prevalence estimates and clustering

Over the course of the summer of 2021, 2063 deer ticks and 678 lone star ticks were sampled from 71 sites (Figure S1). Of these, 161 deer ticks and 2 lone star ticks tested positive for DTV, yielding 7.8% and 0.3% infection rates respectively (Figure 1, Table 1). These were not distributed evenly across the island, as only 19 of the 71 sampled sites yielded positive ticks. In fact, 90 of the 161 (56%) positive ticks derived from only 4 collection sites. Clustering analysis using SatScan identified 8 clusters, 6 of which were statistically significant (Figure 2, Table 2). All of the significant clusters derive from 4 collection sites, which were also grouped into 2 significant clusters with a larger radius.

**Figure 1:**
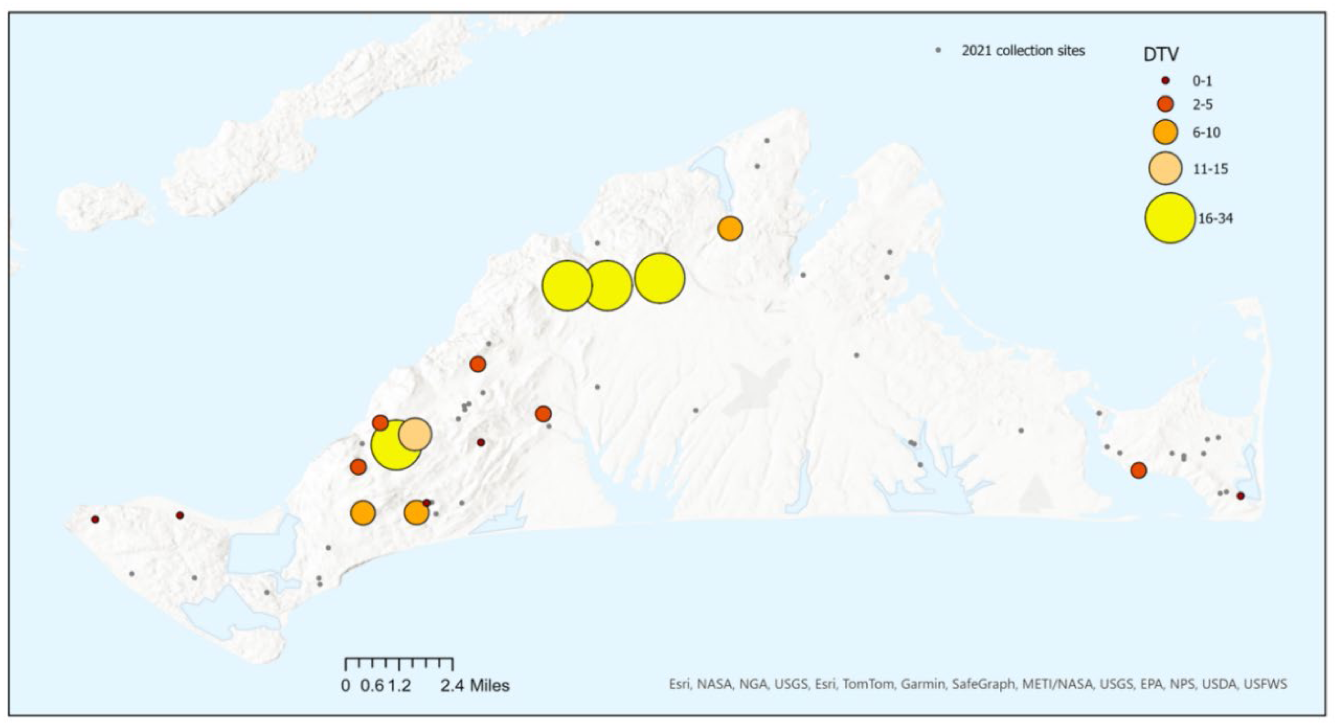
The sites with DTV-positive ticks on MV 2021. Collection sites that yielded no positive ticks are marked with grey dots. Sites with DTV-positive ticks are marked with colored dots that are increasing in size and becoming lighter in color proportional to the number of positive ticks detected.

**Table 1:**
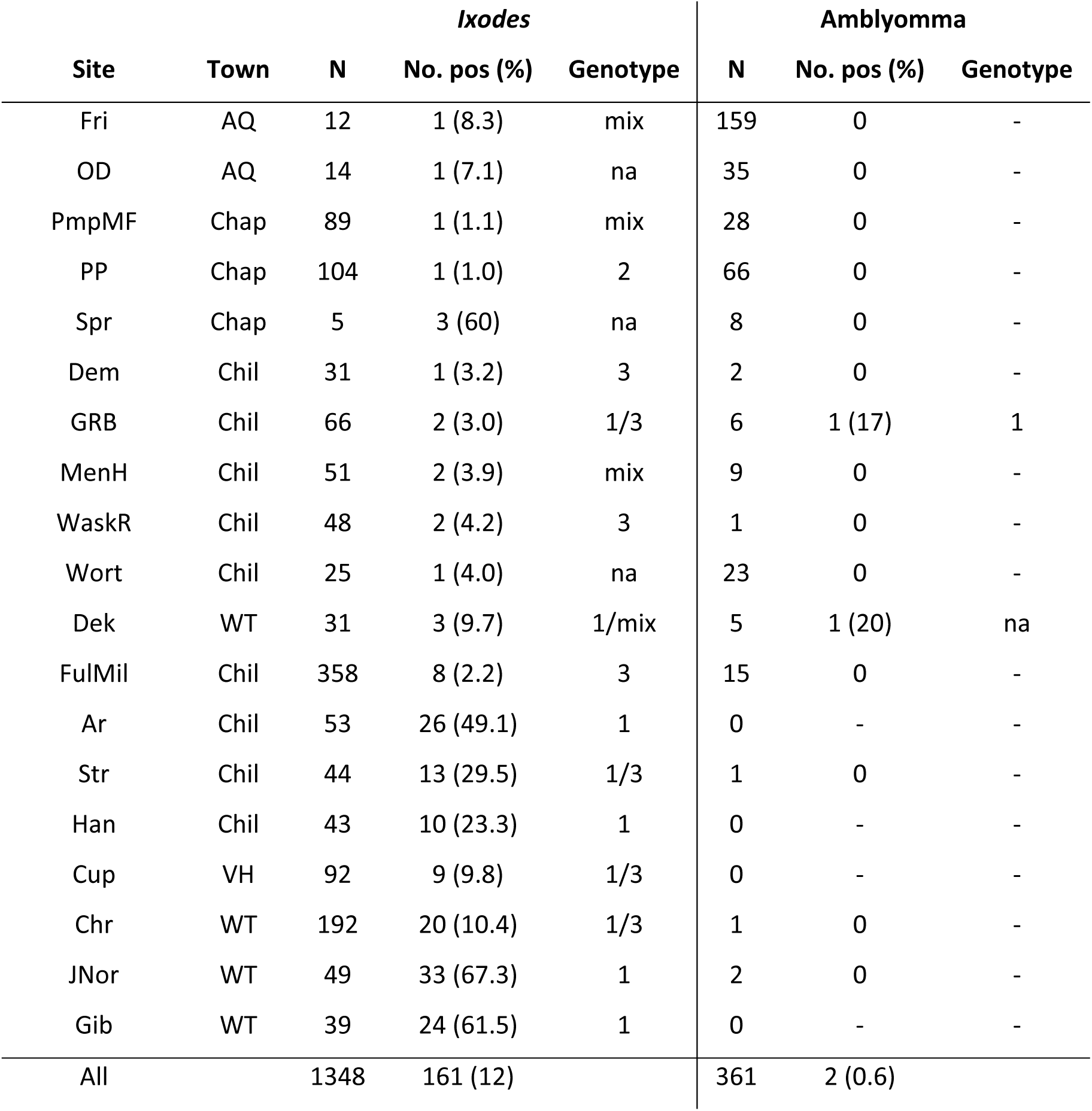
The infection rates of DTV in nymphs from MV 2021 from sites yielding positive ticks. PCR-positive ticks were obtained from 19 of the 71 sites sampled for ticks. Positive ticks were genotyped by sequencing 2 PCR amplicons containing identifying SNPs. Type 1= 1, Type 2=2, Type 3=3, mix= type not identified, likely a mixture, na= no amplification.

**Figure 2:**
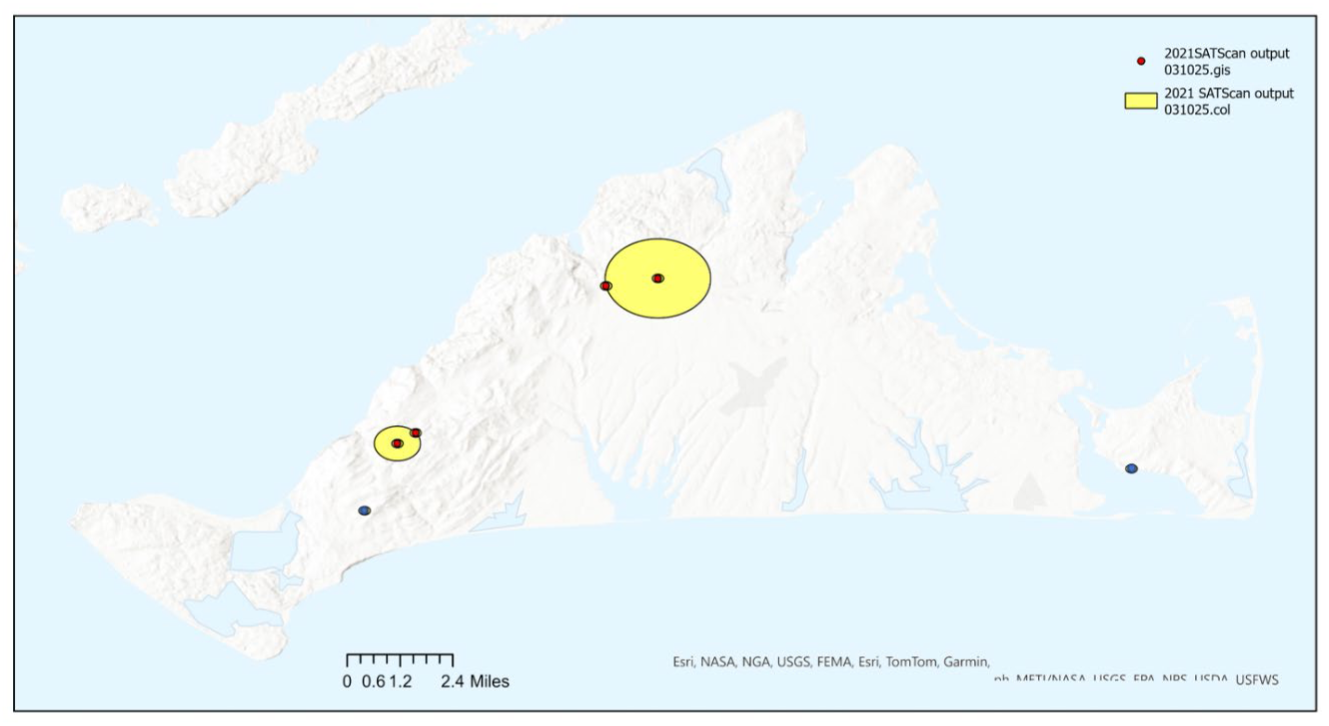
Analysis of clustering of DTV-positive ticks. SatScan was used to identify clusters of DTV-positive ticks. Points indicate that a cluster occurred at a single collection site. Yellow circles indicate the width of a cluster which comprised of more than 1 collection site. Red points indicate statistically significant clusters, and blue indicates clusters that were not statistically significant.

**Table 2:** Cluster analysis of DTV-positive ticks using SatScan. Radius of zero indicates the cluster is from a single collection site.

| Cluster | Site | Radius | No. | No. pos | Relative Risk | p-value |
| --- | --- | --- | --- | --- | --- | --- |
| 1 | JNor-Gib | 1.4km | 88 | 57 | 12.02 | <0.000 |
| 2 | JNor | 0 | 49 | 33 | 10.43 | <0.000 |
| 3 | Ar-Str | 0.63km | 98 | 33 | 6.31 | <0.000 |
| 4 | Gib | 0 | 39 | 24 | 8.96 | <0.000 |
| 5 | Ar | 0 | 54 | 26 | 7.06 | <0.000 |
| 6 | Str | 0 | 44 | 13 | 3.98 | 0.01 |
| 7 | Han | 0 | 43 | 10 | 3.07 | 0.172 |
| 8 | Spr | 0 | 5 | 3 | 7.72 | 0.333 |

### Phylogenetic analysis and genotyping

Whole viral genomes were sequenced in a subset of ticks by amplifying overlapping tiles by PCR and then sequencing using Oxford Nanopore flow cell technology, as was done previously for other viruses (Grubaugh et al., 2019; Quick et al., 2017). Viral genomes were assembled using Geneious Prime, aligned with 2 genomes from GenBank, and a consensus neighbor-joining tree with 500 bootstrap replicates was constructed (Figure 3). This analysis showed that there were 3 genetically distinct types circulating in ticks on MV. Due to the difficulty of successfully obtaining whole viral genomes directly from tick nucleic acid extracts, a new typing assay was developed that relies on SNPs within a single smaller amplicon. The SNPs used to identify each genotype are listed in

**Figure 3:**
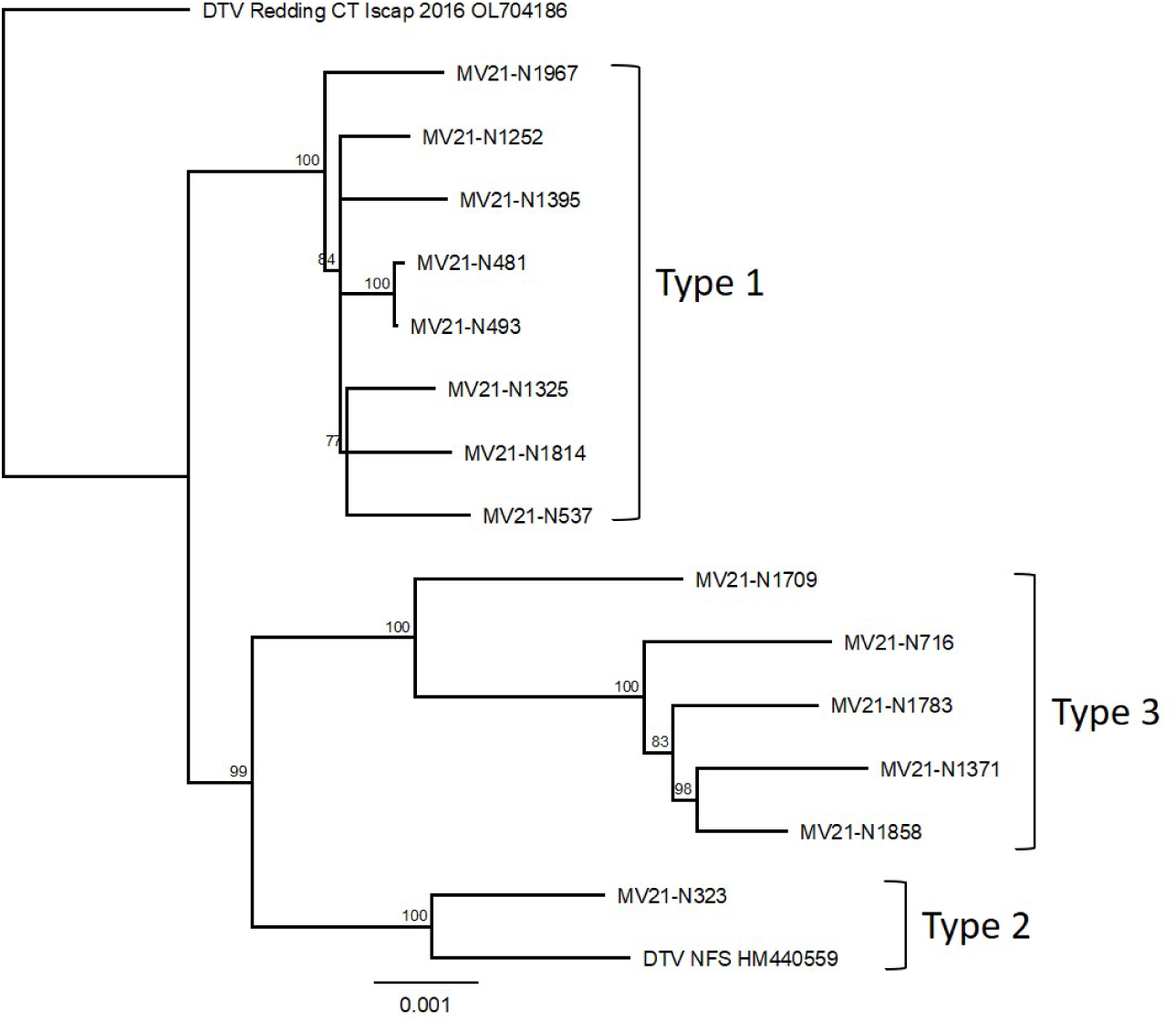
Neighbor-joining consensus tree of DTV genome sequences amplified directly from ticks. The whole genome was sequenced using overlapping PCR-amplified tiles next gen Oxford Nanopore technology. Consensus sequences were assembled using GeneiousPrime. Sequences were aligned to 2 DTV genomes from GenBank, OL704186 from Connecticut and HM440559 from Nantucket, and a consensus tree was constructed using the neighbor-joining algorithm with 500 bootstrap replicates within Geneious TreeBuilder.

Table 3. Of the 161 DTV-positive deer ticks, 94 (58.4%) were determined to be type1, 1 (0.6%) type 2, 22 (13.7%) type 3, 12 (7.5%) contained low numbers of reads with a mixture of sequences that could not be confidently resolved into individual genotypes, and 32 (19.7%) failed to amplify with either set of PCR typing primers. Presumptive virus from one lone star tick nymph was successfully typed to type 1, while the other failed to amplify. Ticks that did not amplify in the typing assay are unlikely to be false positives as we were able to reamplify all positives with a duplex RT-qPCR that amplified DTV along with the GAPDH housekeeping gene. Using a ratio of the crossing threshold for the DTV qPCR (DTV Ct) to that from the GAPDH qPCR (GAPDH Ct), we estimated the relative quantity of virus within the ticks (Figure 4). This analysis demonstrated that the majority of the ticks had low copy numbers of virus (ie high DTV Ct compared to GAPDH); there was no difference between genotype 1 and genotype 3. The single genotype 2 sample had relatively high copy number of virus. Finally, the ticks that did not amplify with the typing primers all had very high Cts, indicative of low levels of virus which likely explains the failure to amplify.

**Table 3.** SNPs used to genotype DTV amplified directly from tick extracts.

| MVamp1 | 5217 | 5223 | 5304 | 5310 | 5313 | 5337 | 5358 | 5382 |
| --- | --- | --- | --- | --- | --- | --- | --- | --- |
| type 1 | G | T | G | G | T | T/A | G/A | T |
| type 2 | G | T | A | A | C | T | G | C |
| type 3 | T | C | A | A | T | T | G | T |
| MVamp2 | 6234 | 6246 | 6255 | 6300 | 6327 | 6330 | 6510 |  |
| type1 | T | C | T | C | T | T | A |  |
| type2 | T | C | C | C | C | T | G |  |
| type3 | C | C | C | T | T | T | A |  |

**Figure 4:**
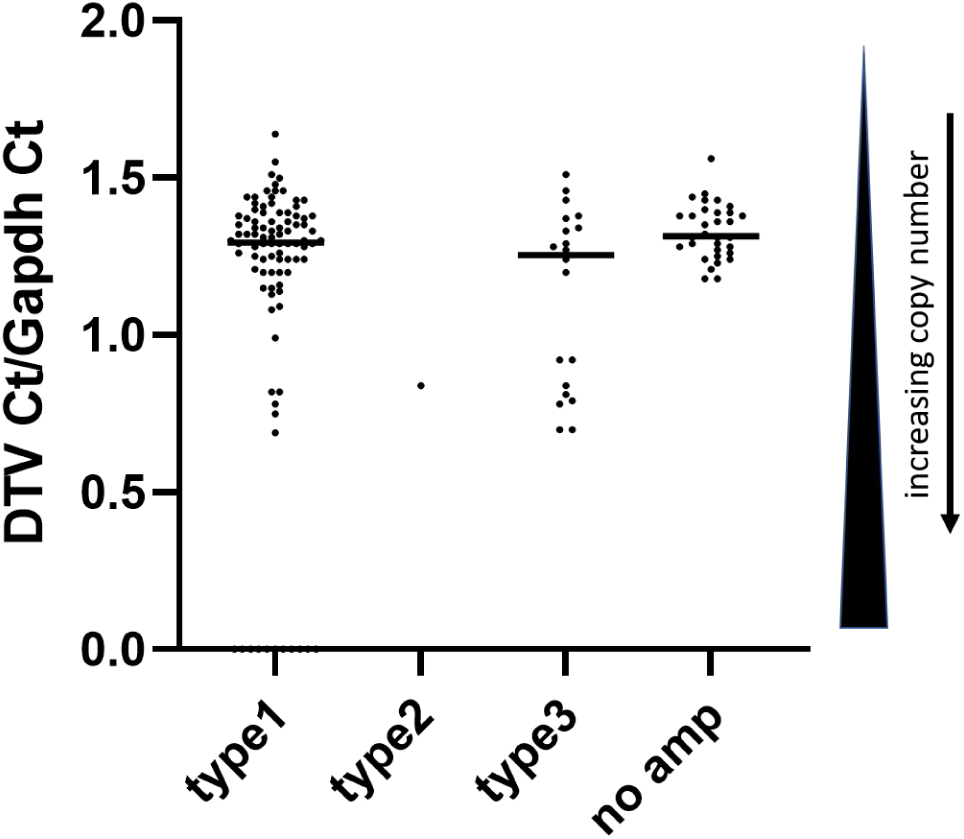
Relative quantitation of DTV in ticks from MV by genotype. DTV-positive ticks were genotyped using SNPs from small 400bp amplicons. The relative amount of virus in each tick is quantitated by running a duplex qPCR with DTV Ns5 primers and *Ixodes* GAPDH primers and dividing the Ct from DTV by the Ct from GAPDH. Note that viremia is inversely related to viremia so that a high DTV Ct is associated with low levels of virus.

Whole genome consensus sequences showed that type 1 and type 3 were 99.28% identical by nucleotides and 99.65% identical amino acids, with 9 differences in their amino acid sequences. These differences were concentrated in the NS2a and NS5 regions of the genome. (Table 4)

**Table 4:** Amino acid differences between DTV type 1 and 3 consensus sequences from this study and sequences from GenBank.

| Gene | AA<br>position | Type 1<br>cons | Type 3<br>cons | NFS<br>HM440559 | CT<br>OL704186 |
| --- | --- | --- | --- | --- | --- |
| Env | 483 | S | N* | S | N* |
| NS1 | 953 | K | K | K | R |
| NS2a | 1179 | V | I | V | V |
| NS2a | 1225 | V | V | A | V |
| NS2a | 1318 | K | N | N | N |
| NS2a | 1351 | F | C | C | C |
| NS3 | 1498 | L | L | S | L |
| NS3 | 1991 | T | T/S | T | T |
| NS4b | 2376 | L | M | L | L |
| NS4b | 2472 | E | E | G | E |
| NS5 | 2784 | K | K | K | R |
| NS5 | 2823 | R | R | R | H |
| NS5 | 2889 | T | A | A | A |
| NS5 | 3161 | K | K | R | K |
| NS5 | 3169 | E | G | G | G |
| NS5 | 3215 | D | D | A | D |
| NS5 | 3391 | K | E | E | E |
| * N associated with neuroinvasive phenotype in mice (Courtney et al., 2025) |  |  |  |  |  |

### Measuring the likelihood of transovarial transmission

To determine whether ticks might have been infected by transovarial transmission, we analyzed the degree of relatedness of PCR-positive ticks. We posit that ticks from a single site that were infected transovarially would derive from the same egg batch (Figure 5). These ticks would have a shared mother and, therefore, share their mitochondrial genome, which is maternally inherited. Accordingly, we sequenced the mitochondrial genome from PCR-positive ticks from yards that contained more than 5 positive ticks and determined whether they were siblings. We did the same for similar number of ticks from control sites and then compared the number ticks with potential siblings for each. Sequence similarity matrixes can be found in the supplemental files. If the ticks were infected by TOT, we would expect that the DTV-positive sites would have significantly more ticks with potential siblings that ticks collected from control sites. In fact, the number of ticks with potential siblings was the similar for both; 34 of 131 (26%) DTV-positive ticks compared to 37 of 132 (28%) control ticks (Table 5). Therefore, we conclude that TOT is unlikely to be contributing significant numbers of infected ticks in our field sites.

**Figure 5:**
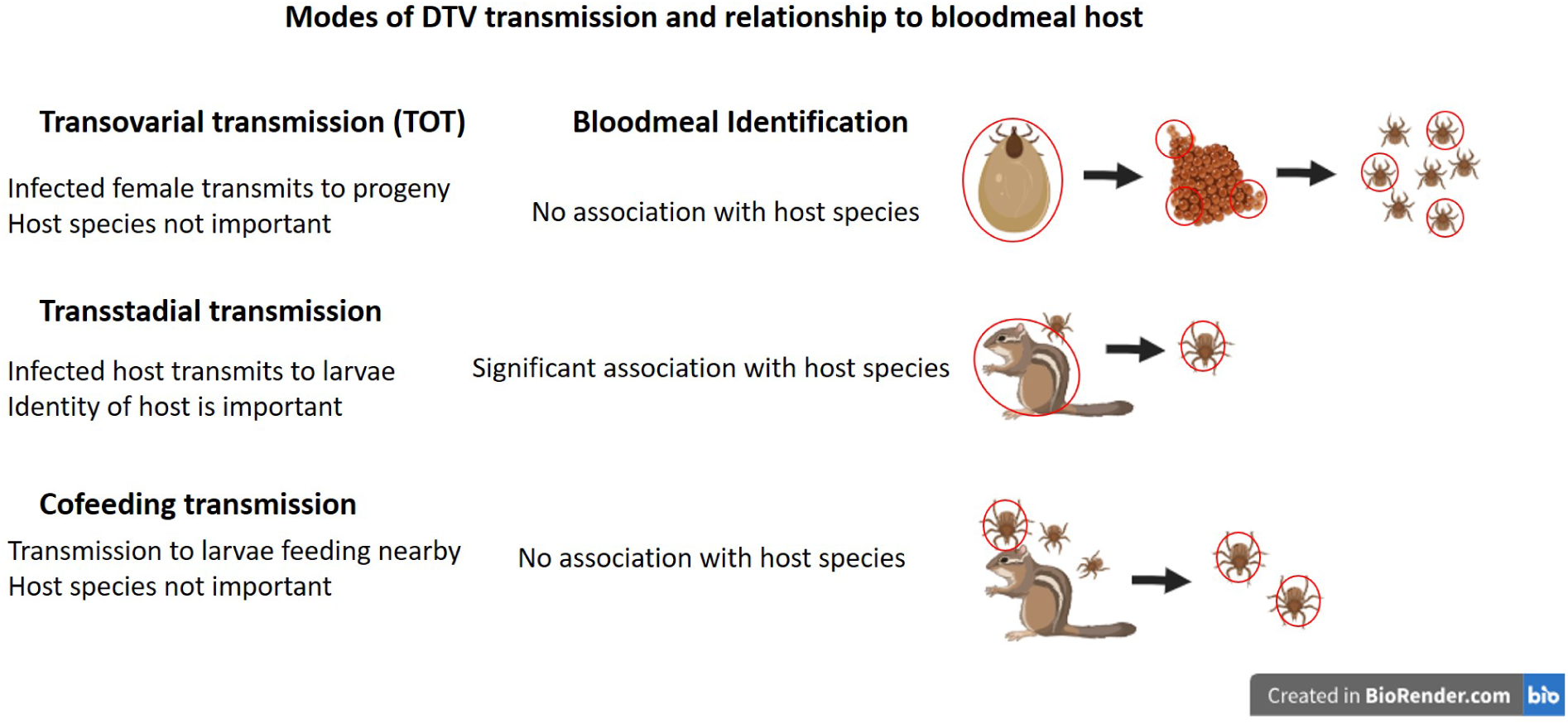
The relationship between mode of transmission and the bloodmeal identification in nymphal ticks.

**Table 5.** Estimation of the contribution of transovarial transmission of DTV at our field sites. Ticks that were collected from the same site and having mitochondrial genomes that are ≥99.95% similar were determined to be potentially from the same egg batch, ie siblings. The number of potential siblings for DTV-infected ticks were compared to controls.

|  | No. tested | No. with a<br>potential sibling | Percent |
| --- | --- | --- | --- |
| DTV-positive | 131 | 34 | 26% |
| Control | 132 | 37 | 28% |

### Investigation into transstadial transmission vs cofeeding

Two other common modes of arboviral transmission are transstadial transmission, in which uninfected larvae acquire infection while feeding on a viremic host and then maintain infection through the molt to emerge as infected nymphs, and cofeeding or nonsystemic transmission, in which larvae can acquire infection by feeding in close proximity to an infected nymph (Figure 5). In the case of transstadial transmission, larvae must feed on a competent/viremic reservoir host. By contrast, with cofeeding the host species does not need to become viremic for transmission to occur and therefore the identity of the host is unimportant (Figure5) (Jones et al., 1987; Labuda et al., 1993). To distinguish between these two modes of arboviral maintenance, we determined the identity of the hosts upon which DTV-positive nymphs had fed on during their larval stage. If transstadial transmission is common, there should be a significant association between infection and host species the tick had fed on. If cofeeding is a common mode of transmission, there should be no such association between host and infection (Figure 5). Bloodmeal identification for DTV-positive ticks determined that shrews had served as the bloodmeal for half (50%) of the infected ticks, and feral cats were the second-most common host, found in 20% (Figure S2).

Positive ticks that had fed on mice, squirrel-chipmunk, rabbit, bird and deer were also found, but each of these contributed fewer than 10% of the ticks. Bloodmeal identification was only successful on one of the DTV-positive lone star ticks, and it was found to have fed on either a raccoon or a skunk (note that this assay cannot differentiate between the two). To determine whether the observed host distribution was indicative of transstadial transmission or cofeeding, we compared the results of bloodmeal analysis for DTV-positive ticks separated by genotype to that for 217 ticks from 9 control sites with no detected DTV-transmission. We found that the hosts identified in ticks with type 3 were not significantly different to that from controls and there were no significant association between DTV infection and any host species. (Figure 6 and 7) By contrast, 53% (50 of 94) of ticks with type 1 had fed on shrews. Compared to bloodmeal hosts from controls, these ticks were significantly more likely to have fed on shrews, and significantly less likely to have fed on squirrels-chipmunks and birds (Figure 6). Ticks that had fed on a shrew were significantly associated with being infected with DTV type 1 (Figure 7). Infection with type 1 was significantly less likely if the tick had fed on a squirrel-chipmunk, rabbit or bird. Finally, the Shannon estimate of diversity for host species identified in ticks with type 1 was significantly lower than that for controls, whereas the diversity estimate for ticks with type 3 did not differ from the controls (Figure 8).

**Figure 6.**
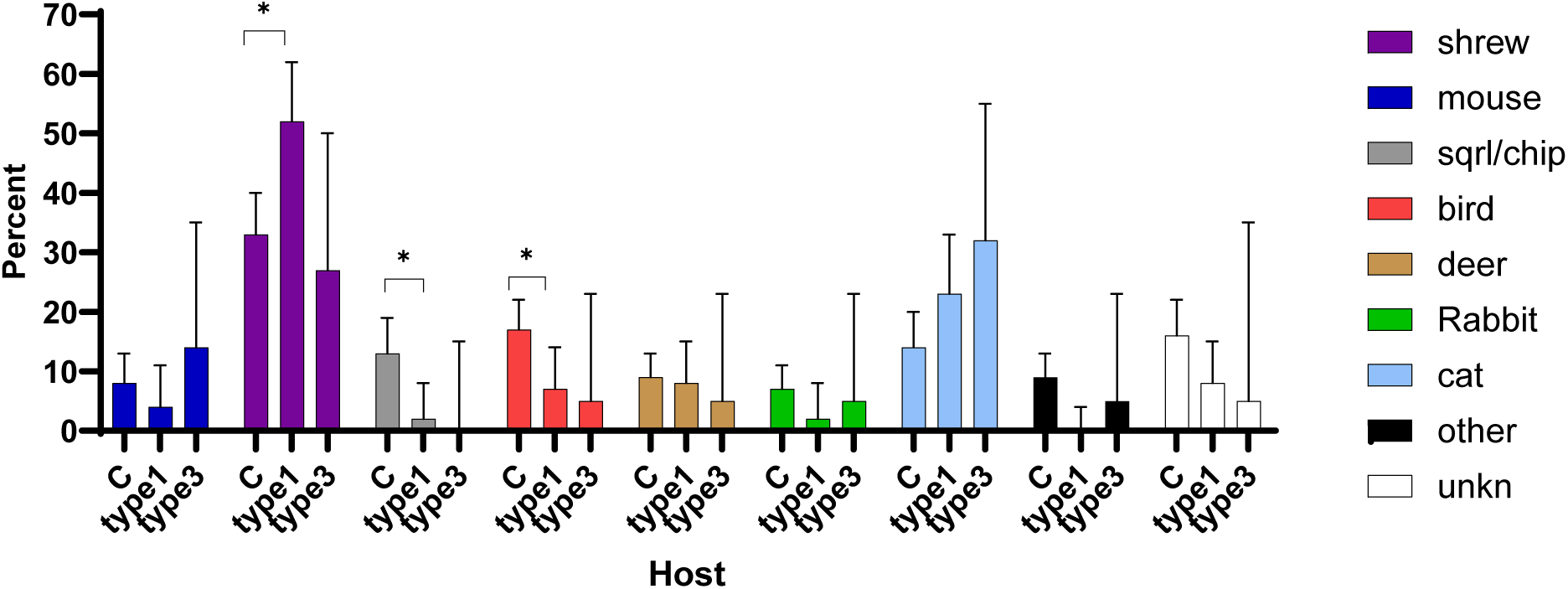
Bloodmeal identification in nymphal deer ticks on MV by DTV infection type. Nymphal ticks were tested for bloodmeal host using PCR to amplify residual mammalian retrotransposons. Bars represent the percentage of ticks that tested positive for each host species with 95% confidence interval. C= ticks collected from control sites, type1= DTV-positive ticks genotype 1, type3= DTV-positive genotype 3, * p <0.05

**Figure 7.**
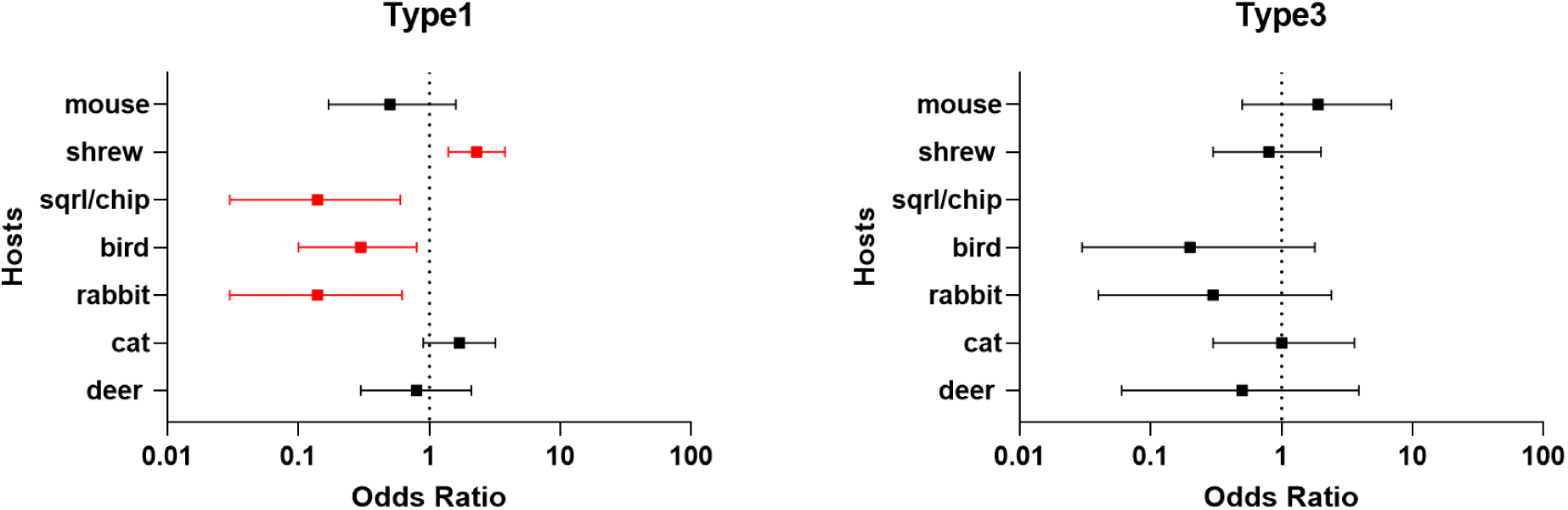
The likelihood that a deer tick infected with either type 1 or type 3 DTV had fed on each host. Odds ratios with 95% confidence intervals were calculated comparing bloodmeal hosts from DTV-positive ticks separated by genotype and to bloodmeal hosts from ticks from control yards that had no evidence of DTV transmission. An odds ratio of 1 is equal to no association; so any confidence interval that cross 1 is not significant. Statistically significant odd ratios are colored in red. No data indicates that there were no DTV ticks testing positive, and the odds ratio was 0 and the 95% confidence interval could not be calculated.

**Figure 8.**
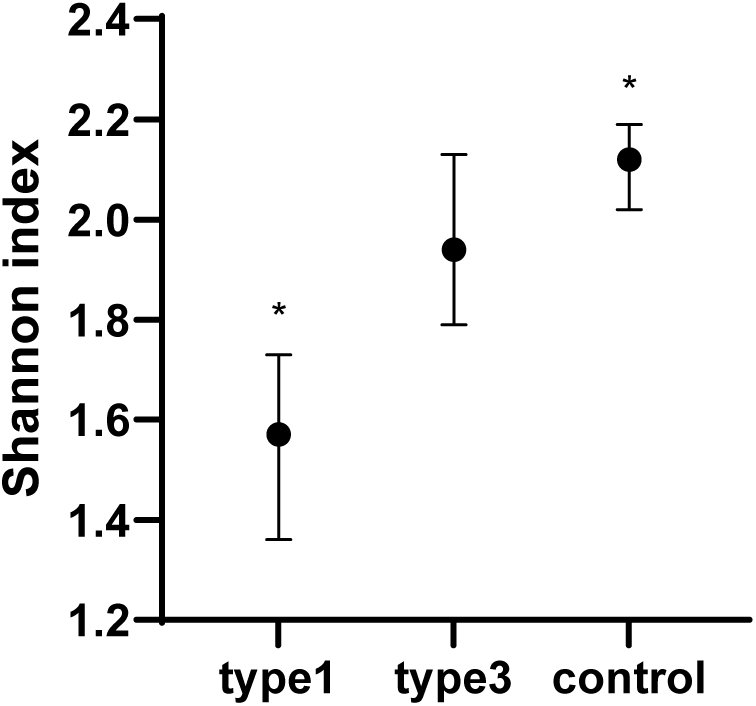
Estimation of the diversity of hosts detected in DTV-positive ticks compared to ticks from control yards. *p<0.05

We conclude that there may be specific associations between DTV type 1 and specific hosts, mainly shrews.

## Discussion

The epizootic of DTV that occurred during the summer of 2021 on MV provided us with the unique opportunity to probe the modes of transmission that serve as the basis for the enzootic cycle of DTV. Prior laboratory and field studies have demonstrated that this virus is capable of being maintained by all 3 major modes of transmission: TOT, transstadial and cofeeding. However, due to the low DTV prevalence typically found in nature, we heretofore have been unable to determine the relative importance of each.

We have been studying the ecology of DTV on MV for more than a decade. Typical infection rates in deer tick nymphs from MV are less than 1%, and therefore the dramatic increase to 7.8% RT-qPCR positive ticks that we observed in ticks collected during 2021 was unexpected. The high Cts detected in the majority of ticks (most over Ct 35, Figure 4) led us to initially suspect PCR contamination. However, we think this is unlikely because the negative controls included in all PCR runs remained clean, and we were able to repeat the positive findings. In addition, a large proportion of positives were confirmed by sequencing, which demonstrated diversity as opposed to uniformity that would be expected for PCR contamination. Although there were some samples which we were unable to confirm by sequencing, this is to be expected due to the difference in sensitivity between the two assays. PCRs that target larger pieces of DNA (cDNA) are inherently less sensitive than qPCR which target smaller pieces. Whether ticks with low viral copy numbers can contribute to the enzootic cycle or comprise a human risk remains to be determined. We did not attempt to culture virus and therefore have no data on the viability of the virus within these ticks. Recent work has identified the presence of defective DTV viral RNA from field-collected ticks (Courtney et al., 2026), and it may be that this is the reason for the low copy number that we detected in these ticks. The issue of host seeking ticks with low viral burdens is worthy of additional study: would they transmit after a longer attachment time, with virus replication during the feeding? or would hosts receiving a low inoculum of virus remain viremic for extended durations due to a reduced immune response?

By sequencing tick mitochondrial genomes, we were able to show that the DTV-positive ticks identified in this study were not any more likely to be related to each other than ticks collected from control yards. This suggests that transovarial transmission is not an important mode of perpetuation on MV but does not completely rule out its contribution to maintenance. Our sampling was extensive across the island but highly focal due to the fact that our collecting efforts were concentrated within residents’ yards. It is possible that nymphs developing from larvae from individual egg batches were distributed more widely than a single yard while attached to the larval host, making it less likely for us to detect infected sibling ticks. The most direct measure of the contribution of TOT to maintenance would be to analyze collections of larvae from our sites, as done in a definitive report of natural POWV TOT (Lange et al., 2024).

Based on our analysis of infected tick relatedness, we assume that the DTV-positive nymphs from this study did not acquire infection transovarially; they must have acquired infection during their larval bloodmeal. This can occur by ingesting virus from either a viremic host or from a cofeeding infected nymph. Previous work with closely-related TBE virus demonstrates that host viremia is not necessary for transmission to occur via cofeeding, and therefore reservoir competence is not necessarily required for maintenance (Jones et al., 1987; Labuda et al., 1993). Transstadial transmission, on the other hand, requires a competent host that becomes viremic, at least transiently. We demonstrated a clear difference between hosts utilized by ticks infected with DTV type 1 and those infected with type 3. Ticks with type 1 were more likely to have fed on shrews than control ticks (Figure 6). Furthermore, having fed on a shrew was significantly associated with infection (Figure 7). Finally, ticks with type 1 showed a low level of species diversity (Figure 8), indicating that these ticks had fed on only a few host species. This type of host association is consistent with a pathogen that is being maintained in ticks largely through transstadial transmission. These data further support our previous work incriminating shrews as a likely reservoir host for DTV (Goethert et al., 2021). By contrast, ticks infected with DTV type 3 did not differ in bloodmeal hosts compared to controls and had no significant associations with any specific host species, consistent with maintenance by cofeeding. We note that fewer ticks with type 3 (22 ticks) were identified compared to type 1 (94 ticks), and this smaller sample size may have decreased the power for finding significant host associations. The odds ratio for a hypothetical association with shrews (the proportion of type 1 ticks identified as having fed on a shrew, 53%) given the smaller number of positive ticks identified as type 3 (11 of 22 type 3 ticks) is 2.4 (95% CI [1.0-5.9] P=0.06). It is likely that if a similarly strong association with a host had occurred for ticks with type 3, we would have had sufficient infected ticks to detect it, but we cannot rule out a weaker association.

The diversity of hosts from ticks infected by DTV type 3 was no different than controls, suggesting that ticks with type 3 had fed on hosts similar to those used by control ticks. We conclude that the two genotypes of DTV on MV are being maintained by different transmission methods.

RNA viruses are known to be highly diverse, due to their reliance on the error-prone RNA-dependent RNA polymerase, often being maintained in what is called a quasispecies (Moya et al., 2004). Our phylogenetic analysis of whole viral genomes shows significant diversity in ticks from a single island. However, our next gen sequencing did not show evidence of quasispecies diversity within the tick, consistent with measurements of inter and intra tick diversity of DTV conducted by Brackney et al. (Brackney et al., 2010). There have been several laboratory-based observations demonstrating phenotypic differences between DTV viral isolates (Courtney et al., 2025; Lange et al., 2025, p. 205; Mcminn et al., 2024). Comparison of consensus sequences showed that there were 9 amino acid differences between type 1 and 3 (Table 4). None of these differences have been described previously, except for S483N found in type 3 which has been associated with a neuroinvasive phenotype (Courtney et al., 2025). It may be that a neuroinvasive virus would have a shorter systemic viremia and be less easily transmitted to feeding ticks, selecting type 3 to depend more on cofeeding ticks for perpetutation. Isolates are required for these two genotypes to test their transmission phenotypes in the laboratory.

The concept of the natural nidality of infection posits that pathogens are often maintained in small foci that are characterized by certain flora and fauna that are crucial to their perpetuation (Pavlovsky, 1966). TBE has long been known to be perpetuated in such foci (Nosek et al., 1970), and by association, DTV is thought to also be similarly maintained. However, the data supporting this conclusion for DTV is sparse, because few long-term ecological studies have been conducted on DTV. DTV persisted within a single site in northwestern Wisconsin over a 10-year period (Brackney et al., 2008), and evidence of a hotspot with greater prevalence of infected adult deer ticks has been documented in coastal Maine (Baxter et al., 2024; Robich et al., 2024). We provide additional evidence for the natural nidality of DTV. We identified 4 of 71 sampled sites as having significant clusters of DTV-positive nymphs, JNor, Gib, Ar and Str. These occurred in close enough proximity to also be grouped together into 2 significant clusters containing 2 sites each, JNor-Gib and Ar-Str. Individually, each of these sites had extraordinarily high infection rates, with over 60% of the ticks at JNor and Gib (33 of 49 at JNor, and 24 of 39 at Gib), and 30% (13 of 44) and 49% (26 of 53) at Str and Ar, respectively.

Together, 56% of all positive ticks were found in these 2 combined clusters. We conclude that although infected ticks were found across the island, the majority were clustered into 2 foci that represented intense enzootic transmission.

Almost all of the ticks from the 4 identified clusters were classified as type 1, and the majority had fed on shrews. It may be that these large numbers of DTV-positive ticks derive from just a few infectious individuals that were capable of transmitting infection to the majority of ticks that fed on them. A small proportion of a host population (20%) may serve as the source for the majority (80%) of infections, known informally as the 20-80 rule (Woolhouse et al., 1997). Accordingly, a few shrews may comprise the reservoirs in our sites. However, the home ranges of shrews are quite small; some estimates range from a mean radius of 13 to 59m (Ingles, 1961; Nosek et al., 1972), which is much smaller than the radius of the clusters estimated by SatScan (Table 2). This suggests that the epizootic we studied is unlikely to be the result of a single highly infectious individual. Our bloodmeal identification does not allow us to identify individual animals; so the number of animals that contributed infected ticks is unknown. Also, it is important to note that while 53% of the ticks with type 1 had fed as larvae on shrews; 47% had fed on other hosts, including feral cats, mice, birds, rabbits, squirrel/chipmunks and deer. We cannot determine whether any of these hosts were acting as actively viremic reservoir hosts or as passive hosts to cofeeding ticks, but it is very likely that that cofeeding also contributes to the perpetuation of type 1.

We would expect that such great prevalence of DTV infection in ticks would translate to an increased hazard for residents living on MV. However, a major outbreak of human cases did not occur that summer, with only a single case documented by the CDC (CDC, 2026). It may be due to the fact that the majority of the ticks we detected had low viral burdens (Figure 4) and the inoculum delivered by such ticks is not great enough to cause infection or cause symptoms severe enough to require a visit to the physician. It is also possible that many of the infected ticks contained defective virus (Courtney et al., 2026). Although the asymptomatic to symptomatic ratio for disease due to Powassan virus remains to be thoroughly studied, none of the 38 subjects submitting ticks to a commercial laboratory reported disease despite the presence of POWV in the ticks that had attached to them (Siegel et al., 2024). Even with the rapid transmission of DTV after attachment (Ebel and Kramer, 2004; Feder et al., 2021), the majority of potentially infectious bites may not lead to disease, which is consistent with the great proportion of subclinical TBEV exposures observed in Europe (Brêchet et al., 2025). We caution that simple “surveillance” (“infection rates”) of host seeking ticks clearly represents a maximum theoretical zoonotic risk and should not be considered a realistic metric for public health intervention.

Lone star ticks are expanding their range in the eastern US and there is greater overlap in distribution with deer ticks and areas of DTV transmission. They have recently invaded MV and now have stable infestations across the island (Johnson et al., 2024). Laboratory studies have confirmed that these ticks are competent vectors for DTV (Sharma et al., 2021), and it is possible that they might become infected by cofeeding, as experimental evidence suggests for the invasive Asian longhorned tick (Obellianne et al., 2024). Due to their infestation intensities and aggressive human biting behavior, lone star ticks might be considered to contribute to DTV risk. Our data does not support their role as secondary vectors for DTV. Despite being collected in our sites with great DTV enzootic transmission, lone star ticks were rarely infected. Only 2 positive ticks were identified over the course of this study.

Therefore, we conclude that these ticks are unlikely to be involved in the perpetuation of DTV on MV, nor do they represent zoonotic risk for DTV. All three stages of these ticks are known to feed mainly on deer, which were not associated with DTV transmission at our sites. We did not conduct bloodmeal identification on the lone star ticks from this study because they did not appear to play an important role in the epizootic. We did, however, examine the DTV-positive ticks for evidence of larval bloodmeal host and found that one had fed on a skunk or raccoon (unfortunately, our assay currently cannot differentiate between the two species); the bloodmeal source for the other tick could not be identified. Although seropositive skunks and raccoons have been detected in the wild (Artsob et al., 1984; McLean et al., 1967), they are unlikely to act as reservoir hosts as they sustain low to undetectable viremia (Kokernot et al., 1969; Nemeth et al., 2021). However, infected larvae were removed from skunks and raccoons in New York, but it is unknown whether they had acquired infection from those hosts or from cofeeding ticks, given that the hosts demonstrated no other evidence of POWV infection (Dupuis II et al., 2013). Unfortunately, data from a single tick in this study cannot determine how that tick had become infected.

In conclusion, the epizootic of DTV on MV during the summer of 2021 allowed us to probe the modes of perpetuation on the island. Whole genome sequences identified 3 distinct genotypes circulating that summer, with the majority of ticks harboring type 1. Bloodmeal remnant analysis of nymphs with type 1 were significantly associated with having fed on a shrew as larvae, suggesting that this genotype is being maintained primarily by transstadial transmission from a competent reservoir host, the shrew. Ticks with type 3, however, demonstrated no specific host associations which is consistent with a pathogen that is being transmitted primarily by cofeeding. The finding that closely-related viral genotypes are perpetuated by different modes of transmission in the same sites speaks to the ecological complexity of arboviral-host-pathogen relationships.

## Supporting information

Supplemental primers

## Acknowledgements

This work was supported by NIH R01 AI 130105 and R01 AI 152209. We have no conflicts of interest to declare.

