## Supplemental primers for "Deer tick virus genotypes are perpetuated by different modes of transmission"

Primers used for screening ticks for DTV and quantitating the relative amount of virus in the ticks.

| Primer name | Sequence | Size | Target | Reference |
| --- | --- | --- | --- | --- |
| POW9466-F1 | ACCATAACAAACATGAAAGTCCAACT | 72bp | DTV NS5 | (El Khoury et al., 2013) |
| POW9466-F2 | CCATCACAAACATGAAAGTCCAACT |  |  |  |
| POW9537-R1 | TGAGTCTGCTGGTCCGATGAC |  |  |  |
| POW9537-R2 | CTGTGAGTCAGCTGGTCCTATGAC |  |  |  |
| POW9453-P | 6FAM-CCTTCCATCATGCGGAT-MGB |  |  |  |
| IscGAPDH-334f | GTG ATC ATC TCC GCA CCT TC | 100bp | Ixodes GAPDH | This study |
| IscGAPDH-433r | ACG AGG CGT TGC TCA CTA TC |  |  |  |
| IscGAPDH-P | HEX-GGGAGTGAACCACAACTCGT-ZENBLHQ |  |  |  |

Tiling primers used to amplify the whole genome of DTV.

| **name** | **pool** | **seq** |
| --- | --- | --- |
| DTV-500_1_LEFT | 1 | GTGTGCGGACACTTTAGTCAGT |
| DTV500-1L-2 | 1 | GTGTGTGGGCACTTTAGTCAGT |
| DTV-500_1_RIGHT | 1 | CGCTAGCTCGCATGACCATAT |
| DTV-500_2_LEFT | 2 | CGTCGACTGGACATGGACTTTC |
| DTV-500-2L-2 | 2 | CGTCGACTGGACATGGRCTTTC |
| DTV-500_2_RIGHT | 2 | ACCCTAGCCATCCAACTATCCA |
| DTV-500_3_LEFT | 1 | AGGGCTGGATGTGGAAAAACAA |
| DTV-500_3_RIGHT | 1 | CTCGCATTCAAACTTGGCACAG |
| DTV-500_4_LEFT | 2 | AACATGGTGTGCAAGAGAGACC |
| DTV-500_4_RIGHT | 2 | AGTCTGATCCCCCAGATTGAAGA |
| DTV-500_5_LEFT | 1 | ACACAAAGACAATCAGGACTGGA |
| DTV-500_5_RIGHT | 1 | CTGGAACCACTGCTGACTTAGG |
| DTV-500_6_LEFT | 2 | CCCCAACCCAACCATTGAAACA |
| DTV-500_6_RIGHT | 2 | ATAGCCGTCATACCACTCCGAT |
| DTV-500-6R-2 | 2 | ATAGCCGTCATACCACTCCGAY |
| DTV-500_7_LEFT | 1 | GGAGCAGACTATGGATGTGCAG |
| DTV-500_7_RIGHT | 1 | AAAAACACCTTTGTGCGTAGGC |
| DTV-500_8_LEFT | 2 | ATGATGATGGGGGTTGATGGAG |
| DTV-500-8Lalt-2 | 2 | TGGAGTGTGCCTGAAAGTYC |
| DTV-500_8_RIGHT | 2 | GTGGTGCTTCTAACTGAGGCTC |
| DTV-500_9_LEFT | 1 | AAACGAAAGGGCCATGGGAC |
| DTV-500_9_RIGHT | 1 | CATCACAAGGGCCATGATCTCT |
| DTV-500_10_LEFT | 2 | ATACTCATGCTGGGCTTGCTTG |
| DTV-500_10_RIGHT | 2 | GCCAAGGGGATGGTAAAGCTTA |
| DTV-500-10R-2 | 2 | GCCAARGGGATGGTAAAGCTTA |
| DTV-500_11_LEFT | 1 | GTATTCCTCACTGTGGCTCTGG |
| DTV-500_11_RIGHT | 1 | TATTCCGGACCAATGGAAAGCC |
| DTV-500_12_LEFT | 2 | CCATGGGGAACTTGCACTTGA |
| DTV-500_12_RIGHT | 2 | AGATTCAGTTTCCCTGGTTGGC |
| DTV-500_13_LEFT | 1 | TGGAGCTTGGAAGGAAAATGGG |
| DTV-500_13_RIGHT | 1 | ATGAAGAGGTGTTGTCAACGGC |
| DTV-500_14_LEFT | 2 | CGCGGGTTGTTCTGAAAGAGAT |
| DTV-500_14_RIGHT | 2 | AGTCTTGCTGTTCAAACAAATCACA |
| DTV-500_15_LEFT | 1 | CGGCTTGGTTCGTGTCATCAAT |
| DTV-500_15_RIGHT | 1 | TCCTTTTCTCCTCCGTCAAACG |
| DTV-500-15R-2 | 1 | TCCTTTTCTCCTCCGTCAARCG |
| DTV-500_16_LEFT | 2 | GGACCAGTTGCCACCTTCTATG |
| DTV-500_16_RIGHT | 2 | GTCCCTCTCTGCCATCTTCATG |
| DTV-500_17_LEFT | 1 | TACCGGACTTGTTGCGCTTG |
| DTV-500-17L-2 | 1 | TACCGGACYTGTTGCGCTTG |
| DTV-500_17_RIGHT | 1 | TTTGTCCATGTGTCCCACACTC |
| DTV-500_18_LEFT | 2 | GGCGGCTAATGAATTGGGCTAT |
| DTV-500_18_RIGHT | 2 | CTTTGGCTCTCCATCGGGAATG |
| DTV-500_19_LEFT | 1 | CTGGGCTAGAGGCTGAATTGAC |
| DTV-500_19_RIGHT | 1 | TCTCGGTTGGTCTCCATGACTC |
| DTV-500_20_LEFT | 2 | ACATCTGGAAGCAACGGTTGAA |
| DTV-500_20_RIGHT | 2 | TGCTCCATCAAAAGGATCACCC |
| DTV-500_21_LEFT | 1 | TACAATCCTGTGTGACATCGGG |
| DTV-500_21_RIGHT | 1 | TACTGCCATGTTCTGTAGGGGT |
| DTV-500_22_LEFT | 2 | CTAGTGATGTGATGGAGCGCAT |
| DTV-500_22_RIGHT | 2 | TGTCTAGAGCGCTCTTCATCCA |
| DTV-500_23_LEFT | 1 | CGCATGGTCGGATGAACAGAAT |
| DTV-500_23_RIGHT | 1 | AACCACTTTTGCGTGATAGGCT |
| DTV-500_24_LEFT | 2 | CAACGCAGACTTGGAGGATGAA |
| DTV-500_24_RIGHT | 2 | TGGAAATGGTGTGAGCAGAAGG |
| DTV-500_25_LEFT | 1 | GAATGACATGGCAAAAACCCGT |
| DTV-500_25_RIGHT | 1 | GTGAGGAACATACCAGATCCTGTG |
| DTV-500_26_LEFT | 2 | GGTCTGGATTTTGGACAACCCT |
| DTV-500_26_RIGHT | 2 | TTTTCTCTGACCTGACTGCGTC |
| DTV-500-26R-2 | 2 | TTTTYTCTGACCTRACTGCGTC |
| DTV-500_27_LEFT | 1 | ATGATCTGCACTGGGAGCTCAA |
| DTV-500_27_RIGHT | 1 | ACACCATCTCCTTGTCAGGCTA |
| DTV-500-27R-2 | 1 | CGGAAAAATCCCGGGGAAGA |

Tiling primers used to amplify deer tick mitochondiral genome

| **name** | **pool** | **seq** |
| --- | --- | --- |
| tickmit_1_LEFT | 1 | GTGTTTGAAAATTAAAGTGGCAGAAAGT |
| tickmit_1_RIGHT | 1 | ATCAATTCACTCCAATGGGGCC |
| tickmit_2_LEFT | 2 | CAACTGAAAATCCCCCTCAAATTCAG |
| tickmit_2_LEFTmod | 2 | CAACTGAAAATCCCCCTCAAATTCAR |
| tickmit 2Lalt - 10436R | 2 | CCTAGAGGGTTGCTTGATCCT |
| tickmit_2_RIGHT | 2 | GGAGCCTCTACATGAGCTTTCG |
| tickmit_2_RIGHTmod | 2 | GGAGCCTCTACATGAGCTTTYGG |
| Tickmit-2Ralt- 8356F | 2 | ACAGAAGCTAAAATTATTGAACCTGA |
| tickmit_3_LEFT | 1 | CCTGAGCGTTTACAAGCAGGTA |
| tickmit_3_RIGHT | 1 | AAACTAGCTAAACCCATACAACCTCT |
| tickmit_4_LEFT | 2 | TTGCTTTTTCCACTCTTAGACAATTAGG |
| Tickmit-4Lalt 6713R | 2 | TGTGCTGGATTAGTGATTCACA |
| tickmit_4_RIGHT | 2 | TCTGGAATTTCAATTTCTTGATCACATCA |
| Tickmit4Ralt - 4726 F | 2 | TGGAGCTATATGACCCCCRT |
| tickmit_5_LEFT | 1 | CATGAATTCCGTGAAAACCTGTTGT |
| tickmit_5_RIGHT | 1 | CGCTATACCATCACTACGTCTTCTC |
| tickmit_6_LEFT | 2 | TCGGTTATCAACATCAAGTAAACGGA |
| Tickmit 6Lalt - 3300 R | 2 | GACGTCCAGGAACAGCATCT |
| tickmit_6_RIGHT | 2 | CAACTTCAGCCATTTTACCGCG |
| Tickmit 6Ralt - 1329 F | 2 | TCGGACTGAATTAGGTCAACC |
| tickmit_7_LEFT | 1 | AGCTATGTCTGGTGCCCCTAAT |
| tickmit_7_RIGHT | 1 | CATTCCTATGCGTCTTCTTTATGTATGT |
| tickmit_8_LEFT | 2 | CCTATCAAGATACTCCTTTACTCAGGC |
| Tickmit 8Lalt - 14216 R | 2 | AGAGATGACCAGTTTACTTCACA |
| tickmit_8_RIGHT | 2 | AAAAATTCATAGGGTCTTCTTGTCCC |

Primers used to amplify smaller fragments of DTV for genotyping

| Primer | Sequence | Size | Genome position |
| --- | --- | --- | --- |
| DTVMVamp1- 11 F | CATCAGCAGCATCGCTCAAG |  | 5083 |
| DTVMVamp1 - 347 R | GGCGATTGACATAAGTGGCG | 337bp |  |
| DTVMVamp1 - 440 R | GGTGACCACGTGCTGCTATA | 430bp |  |
| DTVMVamp2 - 7 F | GTGACTTCACACCCTGGCTT | 346bp | 6226 |
| DTVMVamp2 - 352 R | CCTGGGGTCTCATTGAGCAA |  |  |
