## Supplemental figures for "Deer tick virus genotypes are perpetuated by different modes of transmission"

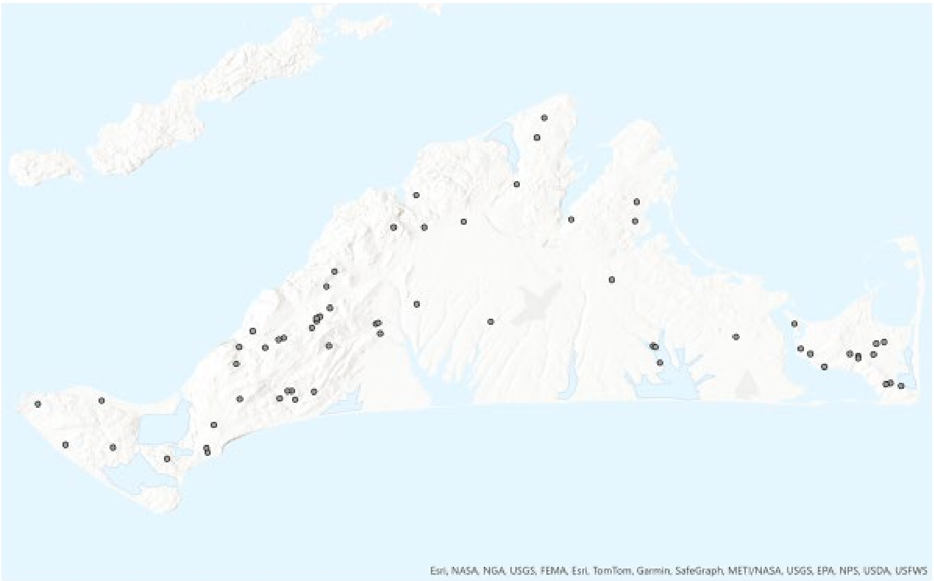


Figure S1: Sampling sites on MV during 2021

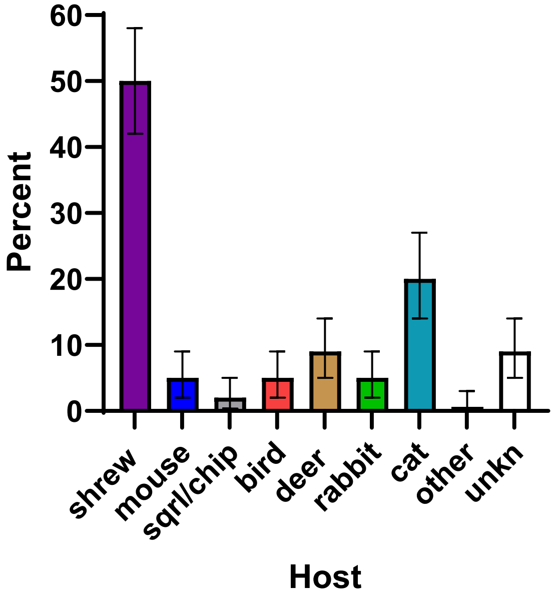


Figure S2. Bloodmeal analysis of deer ticks testing positive for DTV. Bars represent the percentage of DTV-positive ticks that tested positive for each host species.
